# Virus-like particle displaying SARS-CoV-2 receptor binding domain elicits neutralizing antibodies and is protective in a challenge model

**DOI:** 10.1101/2022.11.29.518404

**Authors:** Julia L. McKechnie, Brooke Fiala, Clancey Wolf, Daniel Ellis, Douglas Holtzman, Andrew Feldhaus

## Abstract

While the effort to vaccinate people against severe acute respiratory syndrome coronavirus 2 (SARS-CoV-2) has largely been successful, particularly in the developed world, the rise of new variants as well as waning immunity illustrate the need for a new generation of vaccines that provide broader and/or more durable protection against infection and severe disease. Here we describe the generation and characterization of IVX-411, a computationally designed, two-component virus-like particle (VLP) displaying the ancestral SARS-CoV-2 receptor binding domain (RBD) on its surface. Immunization of mice with IVX-411 generates neutralizing antibodies against the ancestral strain as well as three variants of concern. Neutralizing antibody titers elicited by IVX-411 are durable and significantly higher than those elicited by immunization with soluble RBD and spike antigens. Furthermore, immunization with IVX-411 is shown to be protective in a Syrian Golden hamster challenge model using two different strains of SARS-CoV-2. Overall, these studies demonstrate that IVX-411 is highly immunogenic and capable of eliciting broad, protective immunity.

## Background

The coronavirus disease 2019 (COVID-19) pandemic has led to over 637 million confirmed cases and 6.6 million deaths worldwide as of November 29, 2022 [1]. Our understanding of this disease as well as the causative virus, SARS-CoV-2, has grown dramatically since December, 2019 when it emerged in Wuhan, China [2,3]. Acute SARS-CoV-2 infection is characterized by symptoms such as fever, cough, and sore throat. The virus can also lead to a potentially debilitating condition called ‘long-haul COVID’ [4]. Additionally, the emergence of variants of concern (VOCs) has illustrated the ability of SARS-CoV-2 to evolve, leading to increased transmissibility and immune evasion. Given continued viral evolution and transmission, further advances in SARS-CoV-2 vaccination approaches are necessary to stem the public health impact.

The SARS-CoV-2 spike (S) protein has been the primary target for vaccine development due to the ability of anti-S protein antibodies to neutralize the virus and protect against severe disease [5–9]. S protein trimers decorate the surface of the virion and are comprised of two subunits: S1 and S2 [10,11]. S1 sits at the apex of the S protein and contains the receptor binding domain (RBD). The RBD binds to angiotensin-converting enzyme 2 (ACE2), a protein expressed in human airway epithelia as well as lung parenchyma [12]. After S1 binds ACE2, the S2 subunit facilitates viral fusion with the host cell. During this process, the S protein trimer undergoes a structural transition between its prefusion conformation and its post-fusion conformation, bringing together the fusion peptide and the host cell membrane, mediating virion entry. Antibodies which block S1 binding to ACE2 are often potently neutralizing. Consequently, it is unsurprising that the RBD is the primary target of neutralizing antibodies in human serum [13,14].

Rapid development of vaccines against SARS-CoV-2 was facilitated by two vaccine platform technologies: messenger RNA (mRNA) and viral vectors. While the deployment of these vaccines was instrumental in slowing viral transmission and saving lives, the associated reactogenicity and durability of these vaccines leave room for improvement. Furthermore, the two viral vector vaccines, Ad26.COV2.S (Janssen) and ChAdOX1-S (AstraZeneca), have been linked to a rare blood clotting disorder, thrombosis with thrombocytopenia syndrome [15].

Concern over this potentially life-threatening side effect has led to limited use of these vaccines in developed countries. Waning antibody responses to vaccination, particularly with the mRNA-based vaccines, have also been described [16]. While two doses of the mRNA-based vaccines, BNT162b2 (Pfizer) and mRNA1273 (Moderna), were roughly 90% efficacious in preventing COVID-19 up to two months post immunization, waning antibody responses have resulted in recommendations to receive multiple additional doses [17–20]. Critically, these first generation vaccines are also less efficacious against the omicron variant, which has led to an increase in breakthrough infections [21–23]. Considering these limitations, a new generation of SARS-CoV-2 vaccines that generate broad, durable immunity with reduced reactogenicity are needed.

Virus-like particles (VLPs) are a compelling technology for developing new SARS-CoV-2 vaccines. Whereas soluble recombinant viral proteins tend to be poorly immunogenic, particularly in the absence of an adjuvant, VLPs can induce both humoral and cellular immune responses even without adjuvants [24,25]. One advantage of VLPs is the multivalent presentation of antigen, which promotes B cell receptor clustering and activation, facilitating the production of high affinity antibodies [26]. Importantly, VLP-based vaccines displaying native viral antigens are already commercially available for the prevention of hepatitis B (HBV) and human papillomavirus (HPV) infection. These vaccines have excellent safety and durability profiles [27,28]. Vaccination with HBV vaccines generates high antibody titers that are protective for up to 30 years [29]. Similarly, a single dose of the bivalent HPV vaccine elicits antibody titers that are maintained for at least seven years [30].

Recent advances in computational protein design have allowed for the generation of novel, self-assembling VLPs that can display diverse antigens [24,25,31–36]. Here we produced and further characterized a two-component, computationally designed VLP displaying 60 copies of the ancestral SARS-CoV-2 RBD protein [24], referred to here as IVX-411. We show that immunization of naïve animals with IVX-411 elicits high neutralizing antibody titers against the ancestral strain as well as three VOCs. To evaluate the durability of this immune response, as well as the benefit of the VLP platform over soluble protein, we immunized mice with IVX-411 or soluble S protein. We found that immunization with IVX-411 generated higher neutralizing titers and increased antigen-specific long-lived plasma cells (LLPCs) compared to immunization with soluble protein. Finally, Syrian Golden hamsters immunized with IVX-411 and challenged with SARS-CoV-2 had lower viral loads and reduced disease severity than unimmunized hamsters.

## Methods

### Component production and characterization

RBD-CompA gene based on previously described amino acid sequence [24] was synthesized and cloned by Genscript in the pcDNA3.4+ vector.

DNA was transiently transfected into HEK293F cells, which were incubated at 36 °C with 150 rpm shaking for 4 days before harvest by centrifugation, and 0.2 μm filtration. Ni^2+^ resin (Indigo, Cube Biotech, #75110) was added to 4 μL/mL of cellular supernatant following addition of 1 M Tris pH 8.0 to 50 mM and 5 M NaCl to 300 mM, and incubated with gentle rocking for 2 hours at room temperature (RT) or 16 hours at 4 °C. The loaded Ni^2+^ resin was applied to gravity columns. The columns were washed with 5 column volumes (CV) of wash buffer (20 mM Tris pH 8.0, 300 mM NaCl, 30 mM imidazole, 0.75% CHAPS). Proteins were eluted with elution buffer (20 mM Tris pH 8.0, 300 mM NaCl, 500 mM imidazole, 0.75% CHAPS) and dialyzed into 20 mM Tris pH 8.0, 250 mM NaCl, 5% glycerol, 0.75% CHAPS buffer 3X. The purified RBD-CompA was analyzed by SDS-PAGE and UV-Vis (**Supplemental Figure 1**), and endotoxin levels were determined (Endosafe nexgen-PTS, Charles River; passing value = <10 EU/mg). For manufacture of the CompB pentamer, a transformed *E. coli* Master Cell Bank was expanded into a stirred-tank bioreactor for fed-batch production. CompB was purified using a two-column chromatography process and final formulation conducted by tangential flow filtration. Following purification and formulation, CompB was 0.2 mm filtered and stored at <-65°C.

### VLP production

RBD01-CompA and CompB were quantified by UV-Vis prior to mixing in a 1.2X over equimolar ratio. RBD-CompA was added to a tube to a final concentration of 10 μM, then 20 mM Tris pH 8.0, 250 mM NaCl, 5% glycerol, 0.75% CHAPS buffer was added to bring the reaction volume up to 1 mL. CompB was added to a final concentration of 8 μM. The reaction was mixed and incubated for 1 hour. The resulting VLP was purified by SEC (Superose 6 Increase, Cytiva, #29091596), eluting around 11 mL, using 20 mM Tris pH 8.0, 250 mM NaCl, 5% glycerol, 0.75% CHAPS buffer as the mobile phase. Peak fractions centered around 11 mL were pooled and filter-sterilized (0.2 μm) prior to analysis. VLP concentrations were quantified by UV-Vis.

### UV-Vis spectroscopy

All protein samples were analyzed by UV-Vis on an Agilent Cary 60. Wavelength scans from 400 to 200 nm were collected, with baseline correction using a matching buffer blank. Absorbance at 280 nm was used to quantify the protein concentrations with the molar extinction coefficients and molecular weights as in the following formula:

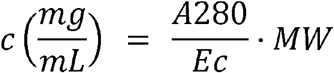

### Dynamic Light Scattering

DLS measurements were taken on a nanoDSF instrument (UNcle, UNchained Laboratories), using autoattenuation of the laser and collecting 10 acquisitions of 5 seconds each at 20 °C.

### Size Exclusion Chromatography

A Superose 6 Increase 10/300 GL column (Cytiva, #29091596) was used to purify trimeric RBD-CompA component, IVX-411 *in vitro* assembly reaction (VLP purification), or IVX-411 purified VLP (analytical), on an AKTA FPLC system (Cytiva, AKTA Go). Columns were equilibrated using 1.2 CV of SEC purification buffer (20 mM Tris pH 8.0, 250 mM NaCl, 100 mM L-Arginine, 5% glycerol, 0.75% CHAPS), then 0.5-2 mL of sample was injected onto the column and eluted using 1.2 CV of SEC purification buffer.

### Biolayer interferometry

Purified RBD-CompA trimeric component, IVX-411 VLP, ACE2-Fc dimerized receptor, and monoclonal antibodies (mAbs) (CR3022, COVA2-39, and CV07-270) were diluted to 10 μg/mL in BLI assay buffer (PBS pH 7.4, 0.5% BSA, 0.05% Tween 20). 200 μL of each dilution and BLI assay buffer were added to black 96-well microplates. Protein A biosensors (Sartorius, #18-5010) were hydrated in BLI assay buffer for 10-20 minutes and loaded onto an Octet Red96 BLI instrument (Pall, FortéBio). Biosensors were dipped into BLI assay buffer to obtain a baseline (60 s), loaded with dimerized ACE2-Fc or mAbs (120 s), dipped into BLI assay buffer, transferred to RBD-CompA and IVX-411 wells (150 s), and dipped back into BLI assay buffer (150 s).

### Negative Stain Transmission Electron Microscopy

IVX-411 sample was diluted to 75 μg/mL in SEC buffer. Sample was adhered to a thick-carbon/Formvar copper 400 mesh grid (Electron Microscopy Sciences, #CF400-Cu-TH) by pipetting 6 μL of sample directly onto the carbon side of the grid and incubating for 1 minute. The grid was dipped into a 50 μL droplet of sterile filtered DI water followed by blotting with grade 1 filter paper (Whatman, #Z240079). The grid was stained by dipping into a 6 μL drop of 0.75% uranyl formate stain, incubated for 1 minute, and blotted off. The staining step was repeated, and the grid dried for 1 minute prior to storage. A Talos L120C TEM microscope, Leginon software, and Gatan camera were used to image the sample.

### VLP Prime-Boost study

The in-life portion of this study was conducted at Abcore Inc. Female, BALB/c mice were immunized IM on days 0 and 21 with 0.2 mg of IVX-411 or IVX-411 + MF59. Immunizations, once prepared at room temperature, were used immediately or within two hours of preparation. Serum samples from each animal were collected on Day 0 (prior to immunization), Day 21 (prior to boost), and on Day 35 (terminal bleed).

### VLP versus soluble protein duration study

The in-life portion of this study was conducted at Aragen Bioscience Inc. BALB/c mice were immunized on days 0 and 21 with 0.2 mg of IVX-411, RBD-CompA, or S-2P formulated with or without MF59. Serum samples were collected on days 0, 20, 35, 63, 91, 119, and 154. Animals were sacrificed on Day 154. Bone marrow was collected for ELISPOT analysis of LLPCs.

### ELISpot

The Mouse IgG ELISpot^BASIC^ kit, Protocol II (Mabtech, #3825-2A) was used by Aragen Bioscience Inc to perform LLPC quantification. SARS-CoV-2 antigen RBD01-dn5B was diluted to 80 mg/mL and used to coat wells of PVDF plates (Mabtech, #3654-WP-10) according to the assay kit protocol. Anti-mouse IgG was diluted to 20 mg/ mL and was used to coat additional wells on the same plate. Plates were washed and blocked with assay media for 30 minutes at RT. Cells isolated from femoral bone marrow were resuspended at 1 × 10^6^ cells/mL. 100 mL was added to the coated wells. Plates were incubated at 33°C for 14-16 hours. Spots counts/well were determined using a Zellnet Consulting ELISpot reader.

### Syrian Golden hamster efficacy study

The in-life portion of this study was conducted at Lovelace Biomedical. 36 male, SGHs were purchased from Charles River. 16 hamsters were immunized IM with 0.2 mg IVX-411 formulated with MF59 on days 0 and 21. Remaining hamsters were immunized with PBS. Serum samples were collected on days 0, 21, and 35. Body weights were measured starting on Day 39. On Day 42 8 IVX-411 immunized hamsters and 8 PBS immunized hamsters were challenged via intranasal instillation with 4.64 × 10^5^ TCID_50_/animal of WA1/2020; 8 IVX-411 immunized hamsters and 8 PBS immunized hamsters were challenged with 1.53 × 10^6^ TCID_50_/animal of B.1.617.2. Animals were sacrificed on Day 47 and lung weights were measured. Lungs were fixed in NBF, trimmed, paraffin embedded, sectioned at 4 mm, and stained with hematoxylin and eosin for microscopic examination. Histopathologic findings were graded subjectively on a scale of 1 to 5. The Provantis™ (Instem LSS Ltd., Staffordshire, England) computer software was used for necropsy and histopathology data acquisition, reporting, and analysis.

### RT-qPCR

RT-qPCR analysis on lung and nasal swab samples collected from the Syrian Golden hamster study were performed at Lovelace Biomedical. Samples were processed in Trizol using a TissueLyser and centrifuged at 4000 x g for 5 minutes. RNA extraction was performed on supernatants using the QIAGEN RNeasy kit according to the manufacturer’s instructions. Samples were run in triplicate and genome copies per mL or gram equivalents were calculated from a standard curve generated from RNA standards of known copy concentration. The N and E gene primers and probe sequences were as follows:

N gene:

SARS-CoV-2 Forward: 5’ TTACAAACATTGGCCGCAAA 3’

SARS-CoV-2 Reverse: 5’ GCGCGACATTCCGAAGAA 3’ SARS-CoV-2 Probe: 6FAM-ACAATTTGCCCCCAGCGCTTCAG-BHQ-1

E gene:

SARS-CoV-2 Forward: 5’ ACAGGTACGTTAATAGTTAATAGCGT 3’

SARS-CoV-2 Reverse: 5’ ATATTGCAGCAGTACGCACACA 3’

SARS-CoV-2 Probe: 6FAM-ACACTAGCCATCCTTACTGCGCTTCG-BHQ-1

### SARS-CoV-2 Pseudo-particle Neutralization Assay (PNA)

PNA assays were performed at Nexelis. ACE-2 expressing Vero E6 cells (ATCC CRL-1586) were seeded in a 96-well microtiter plate at 20,000 cells/well. Serum samples and controls were heat-inactivated at 56°C for 30 minutes, diluted in duplicate in cell medium, and serial two-fold dilutions were performed. Each SARS-CoV-2 pseudovirus was diluted to reach a desired concentration, added to the diluted serum samples, and incubated at 37°C in 5% CO_2_ for 1 hour. This mixture was added to the cells at 80% confluency. Plates were incubated for 18-22 hours at 37°C in 5% CO_2_ before supernatants were removed. 50 mL of ONE-Glo EX Luciferase Assay Substrate (Promega, #E8110) diluted 1:2 in cell media was added to each well and incubated at RT for 3 minutes with agitation. Luminescence across all wavelengths was measured for 0.5 seconds using a SpectraMax iD3 microplate reader and SoftMax Pro v7.0.1 (Molecular Devices). A titration curve using a 4-parameter logistic regression was made for each dilution. The reciprocal dilution of the sample for which the luminescence was equal to a pre-defined cut-point of 50 was reported as the titer. The cut-point was determined using linear regression using 50% flanking points.

## Results

### Production and characterization of two-component I53-50 VLPs displaying ancestral ARS-CoV-2 RBD

A two-component, computationally designed protein VLP, referred to as I53-50 [35], was utilized to display the ancestral (Wuhan-Hu-1) SARS-CoV-2 RBD to improve the immunogenicity of the monomeric antigen as previously described [24]. I53-50 is a 120-subunit VLP comprised of 20 homotrimeric (CompA) and 12 homopentameric (CompB) subunits, capable of *in vitro* assembly following purification and subsequent mixing of the individual components [35,37]. The his-tagged SARS-CoV-2 RBD antigen was displayed on I53-50 by genetically fusing the C-terminus of the antigen to the N-terminus of the CompA subunit using a 16-residue glycine-serine linker (RBD-CompA) [24]. RBD-CompA and CompB were individually expressed in HEK293F and *E. coli* cells, respectively, and purified prior to mixing in equimolar ratios to induce spontaneous self-assembly of VLPs *in vitro* (**Figure 1A**). Following purification by size exclusion chromatography (SEC) to remove residual components, the resulting RBD-I53-50 VLPs (IVX-411) were characterized for identity, aggregation state, antigenicity and VLP structural integrity. UV-Vis spectroscopy wavelength scan analysis showed a peak at 280 nm and low-level scattering typical of non-aggregated, well-formed VLPs (ratio of absorbance at 320 nm to 280 nm = ∼0.1) (**Figure 1B**). Dynamic light scattering (DLS) measurements suggested non-aggregated, monodispersed VLPs with a polydispersity index of 11.8% (**Figure 1C**). Analytical SEC of purified IVX-411 resulted in a resolved peak centered around 11 mL, consistent with the calculated molecular weight of the VLP **(Figure 1D)**. IVX-411 eluted earlier than the constituent components (**Figure 1D**). The antigen appeared intact based on binding to ACE2-Fc, as well as three additional anti-RBD monoclonal antibodies, as measured by biolayer interferometry (BLI) (**Figure 1E**). Finally, negative stain transmission electron microscopy (nsTEM) confirmed that the VLP sample consisted of monodispersed, intact VLPs of the expected diameter (**Figure 1F**).

**Figure 1:**
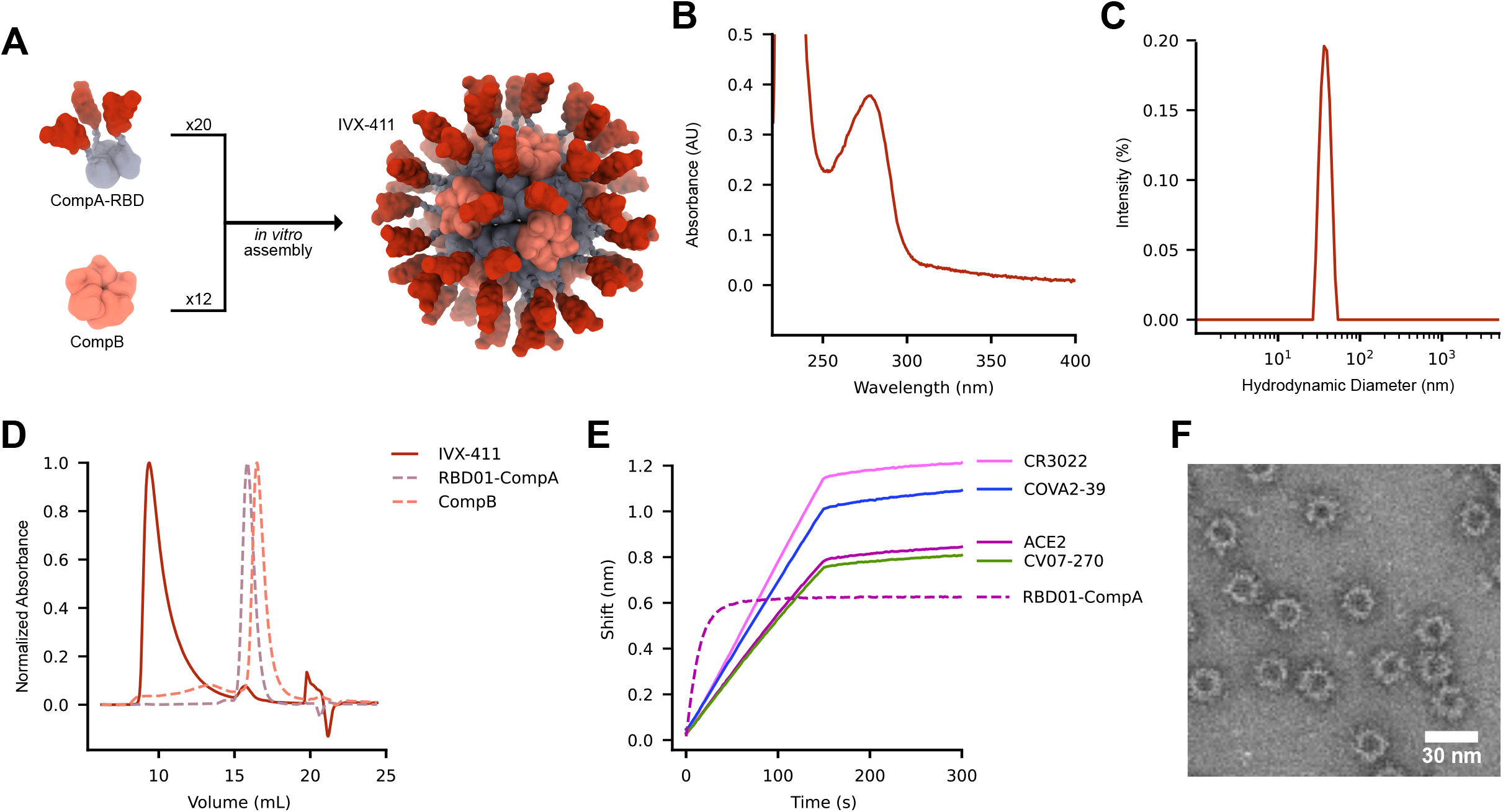
Production and characterization of IVX-411 VLPs. **(A)** Schematic of *in vitro* assembly. **(B)** UV-Vis spectroscopy. **(C)** Dynamic Light Scattering. **(D)** Size Exclusion Chromatography. **(E)** Biolayer Interferometry. **(F)** Negative stain Transmission Electron Microscopy.

### Animals primed and boosted with IVX-411 develop robust neutralizing titers against three VOCs

We evaluated the ability of IVX-411 to induce neutralizing antibody titers against the ancestral SARS-CoV-2 strain as well as three variants of concern (beta, gamma, and delta). Naïve BALB/c mice were immunized intramuscularly (IM) on Day 0 and Day 21 with 0.2 mg of IVX-411 formulated with CSL Seqirus’ proprietary oil-in-water adjuvant, MF59 **(Figure 2A)**. Mice were bled on Day 0 (pre-immunization), Day 21 (pre-boost), and on Day 35 (14 days post-boost). Neutralizing titers against the ancestral strain and the beta variant were measured at all time points using a cell-based pseudo-particle neutralization assay (PNA). Neutralizing titers against gamma and delta variants were measured using only the Day 35 samples.

**Figure 2:**
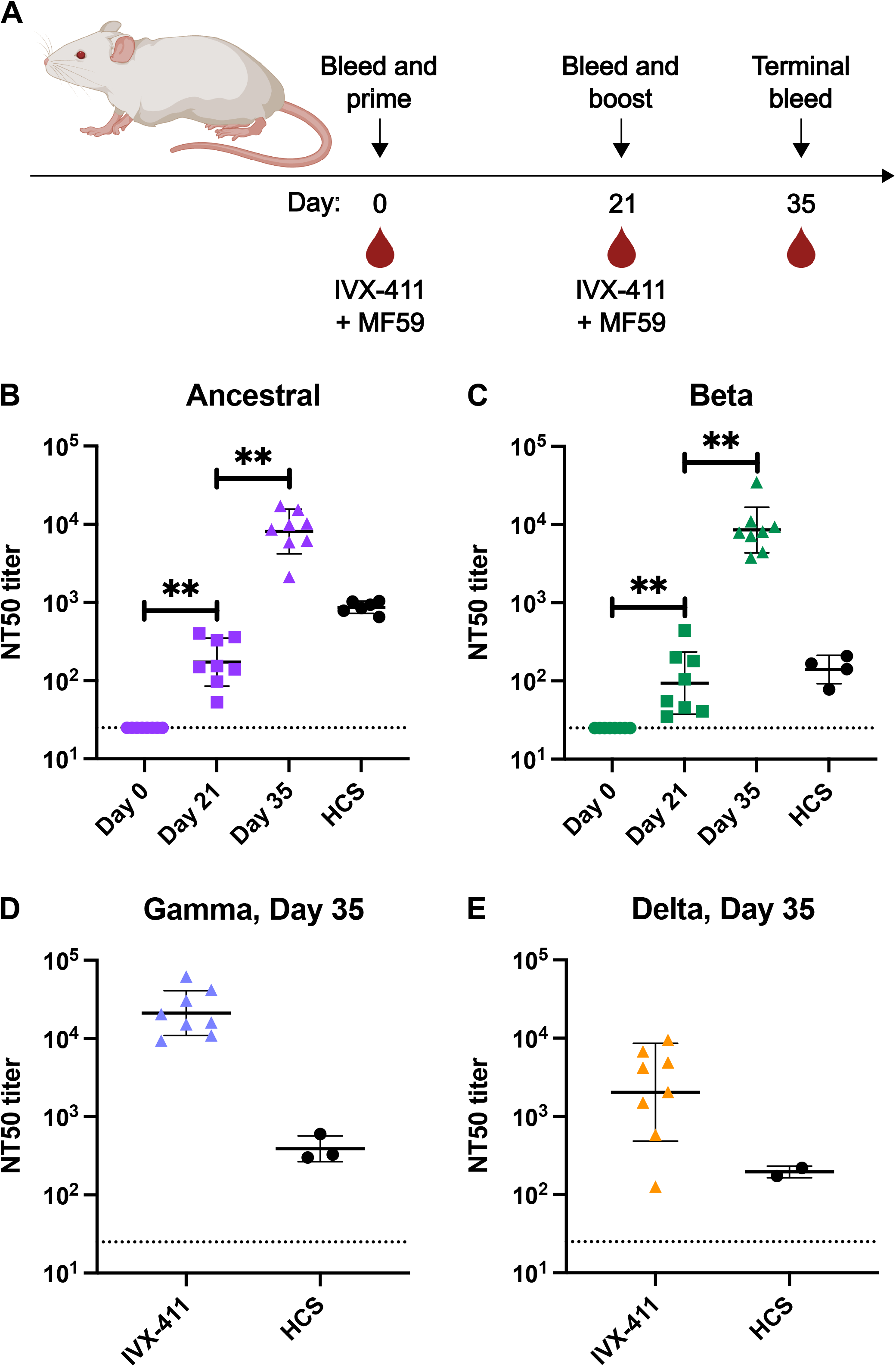
Immunization with IVX-411 elicits neutralizing antibodies against four SARS-CoV-2 strains. **(A)** Female BALB/c mice were immunized IM on Day 0 and Day 21 with IVX-411 formulated with MF59 and bled on Day 0 (pre-immunization), Day 21, and Day 35 (n = 8). Serum neutralizing titers against the ancestral strain **(B)**, as well as the beta **(C)**, gamma **(D)**, and delta **(E)** variants were measured using a cell-based pseudo-particle neutralization assay. Human convalescent serum (HCS) was used as a positive control. Dashed horizontal line represents the lower limit of quantitation. P-values were calculated in GraphPad Prism 9 using a Wilcoxon matched-pairs signed rank test. *, P< 0.05; **, P<0.01.

Three weeks after the initial priming dose, mice immunized with IVX-411 had an average Day 21 neutralizing antibody titer (NT50) of 2.1 × 10^2^ against the ancestral strain **(Figure 2B)** and an average neutralizing antibody titer of 1.4 × 10^2^ against the beta variant **(Figure 2C)**. After receiving a booster immunization, the Day 35 neutralizing titers against the ancestral strain increased 45-fold. Similarly, a 78-fold increase in neutralizing titers against the beta variant was also observed. Mice primed and boosted with IVX-411 had an average neutralizing titer against the gamma variant of 2.58 × 10^4^ **(Figure 2D)**. Neutralizing titers against the delta variant were lower than those against gamma, beta, and the ancestral strain, with an average neutralizing titer of 3.71 × 10^3^ **(Figure 2E)**. Day 35 neutralizing antibody titers against all four strains were higher than those of control human convalescent sera. Together, these results demonstrate that immunization with IVX-411 results in a broad, potent antibody response.

### Immunization with IVX-411 induces a more potent humoral immune response than immunization with soluble protein

Having confirmed the immunogenicity of IVX-411, we next sought to evaluate its ability to elicit durable, humoral immunity compared to trimeric spike-based and RBD-based soluble antigens. Naïve BALB/c mice were immunized on Day 0 and boosted on Day 21 with either 0.2 mg of IVX-411, an equivalent antigen dose of RBD-CompA (0.15 μg), or an equivalent RBD antigen dose of S-2P (0.4 μg), a stabilized prefusion version of the S protein ectodomain **(Figure 3A)**. Each antigen was formulated with either MF59 or an aqueous buffer. Serum samples were collected on days 0, 20, 35, 63, 91, 119, and 154. On Day 154 the animals were sacrificed, at which point LLPCs were isolated from the bone marrow and assessed by ELISpot.

**Figure 3:**
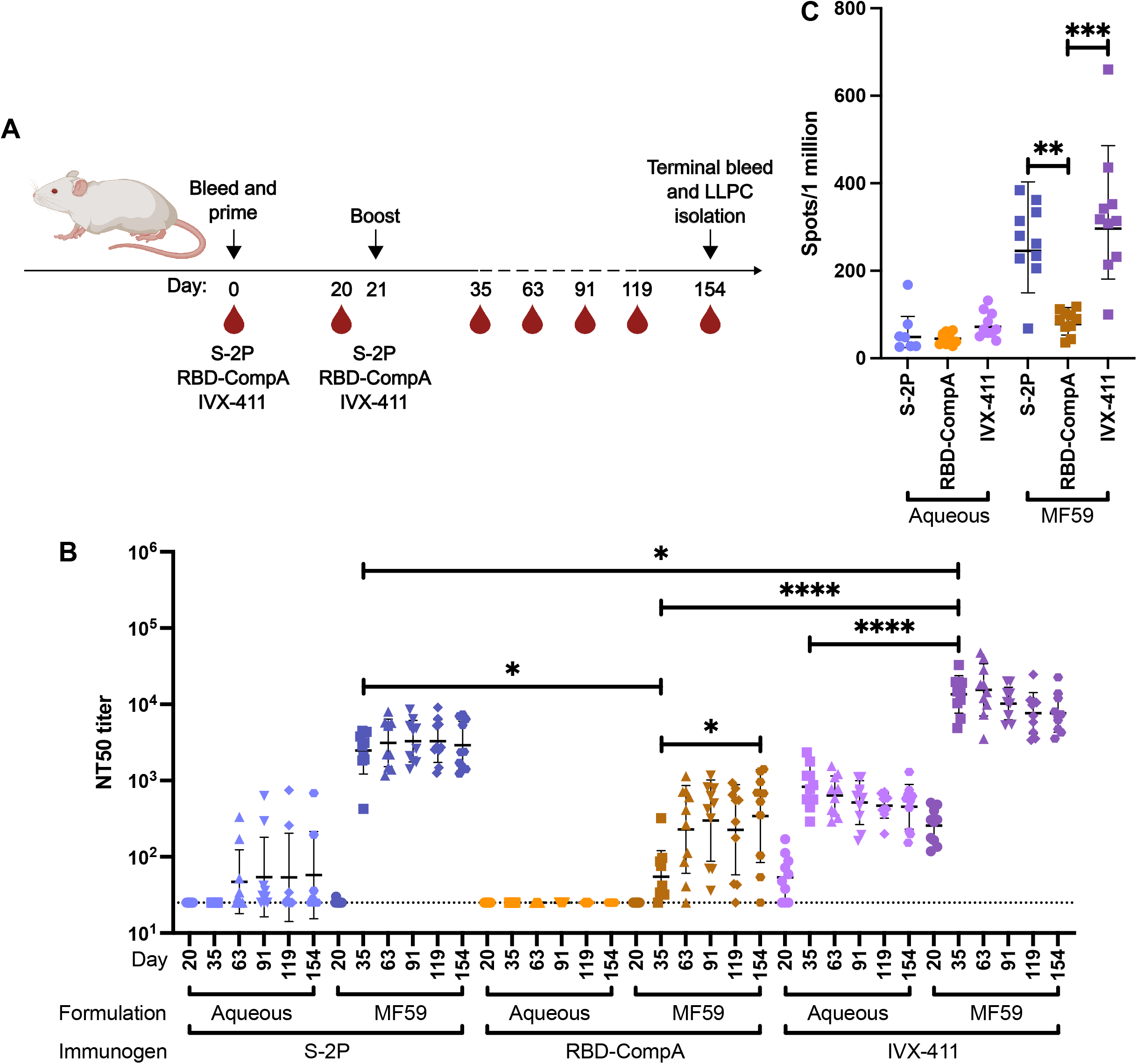
Immunization with IVX-411 elicits higher neutralizing titers and a larger antigen-specific LLPC compartment than immunization with soluble protein. **A)** Female BALB/c mice were immunized IM with IVX-411, RBD-CompA, or S-2P formulated with or without MF59 on Day 0 and Day 21. Serum was collected on days 0, 20, 35, 63, 91, 119, and 154. Mice were sacrificed and bone marrow was collected on Day 154 for isolation of LLPCs (n = 10). **(B)** Serum neutralizing titers against the ancestral SARS-CoV-2 strain. **(C)** ELISpot measuring the frequency of RBD-specific LLPCs in the bone marrow. Dashed horizontal line represents the lower limit of quantitation. P-values were calculated in GraphPad Prism 9 using a Mann-Whitney test or a Kruskal-Wallis test followed by a Dunn’s multiple comparisons test. *, P< 0.05; **, P<0.01; ***, P<0.001; ****, P<0.0001.

Serum analysis by a cell-based PNA revealed that immunization with IVX-411 generated higher neutralizing antibody titers than immunization with soluble proteins. On Day 20, after a single immunization, only animals immunized with IVX-411 had neutralizing antibody titers above the lower limit of quantitation **(Figure 3B)**. Even at this early timepoint, MF59 enhanced the neutralizing antibody response in naïve animals with the IVX-411 MF59 group having 4.4-fold higher Day 20 neutralizing antibody titers compared to animals immunized with IVX-411 alone. This trend was consistent throughout the duration of the study. Compared to the aqueous formulations, immunization with antigen plus MF59 resulted in 16, 3.1, and 116-fold increases in Day 35 neutralizing antibody titers for IVX-411, RBD-CompA, and S-2P, respectively. Importantly, the Day 35 neutralizing antibody titers in the IVX-411 MF59 group were 198-fold higher than those in the RBD-CompA MF59 group and 5.3-fold higher than those in the S-2P MF59 group. These results suggest that immunization with IVX-411 results in higher neutralizing antibody titers than immunization with soluble protein, particularly compared with soluble trimerized RBD, which is poorly immunogenic on its own. There were no statistically significant differences in the neutralizing antibody titers within the different treatment groups from Day 35 to Day 154, except for the RBD-CompA MF59 group. Unlike the other groups, neutralizing antibody titers following a second administration of RBD-CompA MF59 did not reach their peak until Day 154. The 7.9-fold increase in titers on Day 154 compared to Day 35 rose to the level of statistical significance. While the overall consistency of neutralizing antibody titers post-boost reveals the durability of the humoral immune response elicited by immunization in general, these results show that immunization with IVX-411 leads to higher neutralizing titers. To further evaluate the ability of IVX-411 to generate a long-term humoral immune response, an ELISpot assay was performed on isolated LLPCs. Similar to the neutralizing antibody titer results, these results demonstrated adjuvantation was key to generating a sizable RBD-specific LLPC compartment in naïve animals. There were no statistically significant differences in the number of RBD-specific LLPCs between the aqueous conditions. **(Figure 3C)**. Compared to the aqueous formulations, immunization with antigen plus MF59 resulted in 4.3, 1.8, and 4.4-fold increases in LLPC counts for IVX-411, RBD-CompA, and S-2P, respectively.

Immunization with IVX-411 MF59 resulted in a statistically significant, 3.9-fold increase in the number of LLPCs compared to immunization with RBD-CompA MF59. The 1.2-fold increase in LLPC counts observed in the IVX-411 MF59 immunized group compared to the S-2P MF59 immunized group was modest and did not reach the level of statistical significance. These results demonstrate that immunization with IVX-411 in the presence of an oil-in-water emulsion induces durable neutralizing antibody titers, which along with LLPC counts, are superior to those induced by immunization with soluble antigens.

### Immunization with IVX-411 reduces disease severity and viral load in Syrian Golden hamster model

To evaluate whether IVX-411 could protect against SARS-CoV-2 infection, a viral challenge study using WA1/2020 (ancestral) and B.1.671.2 (delta) strains was performed in Syrian Golden hamsters. Hamsters were randomly assigned to five treatment groups: PBS, Unchallenged; PBS, WA1/2020; PBS, B.1.617.2; IVX-411, WA1/2020; and IVX-411, B.1.671.2 **(Figure 4A)**. On Day 0 and Day 21 hamsters in the IVX-411, WA1/2020, and IVX-411, B.1.617.2 groups were immunized with IVX-411 formulated with MF59. Hamsters in the other three treatment groups (PBS, Unchallenged; PBS, WA1/2020; and PBS, B.1.617.2), were injected with PBS. Hamsters were weighed and bled on days 0, 21, and 35. On Day 42 all hamsters, except those in the PBS, Unchallenged group, were inoculated intranasally with either 4.64 × 10^5^ TCID_50_/animal of WA1/2020 or 1.53 × 10^6^ TCID_50_/animal of B.1.617.2. Body weights were recorded daily from Day 39 to Day 47, at which point animals were sacrificed and lung tissue as well as nasal swabs were collected. Hamsters challenged with either WA1/2020 or B.1.617.2 began to lose weight four to five days post challenge **(Figure 4B)**. By six days post challenge, hamsters immunized with IVX-411 had significantly higher body weights compared to their unimmunized counterparts. Interestingly, this reduction in weight loss was more pronounced in animals challenged with B.1.617.2 compared to WA1/2020, despite the higher B.1.617.2 challenge dose. This continued to be the case until the termination of the study on Day 47.

**Figure 4:**
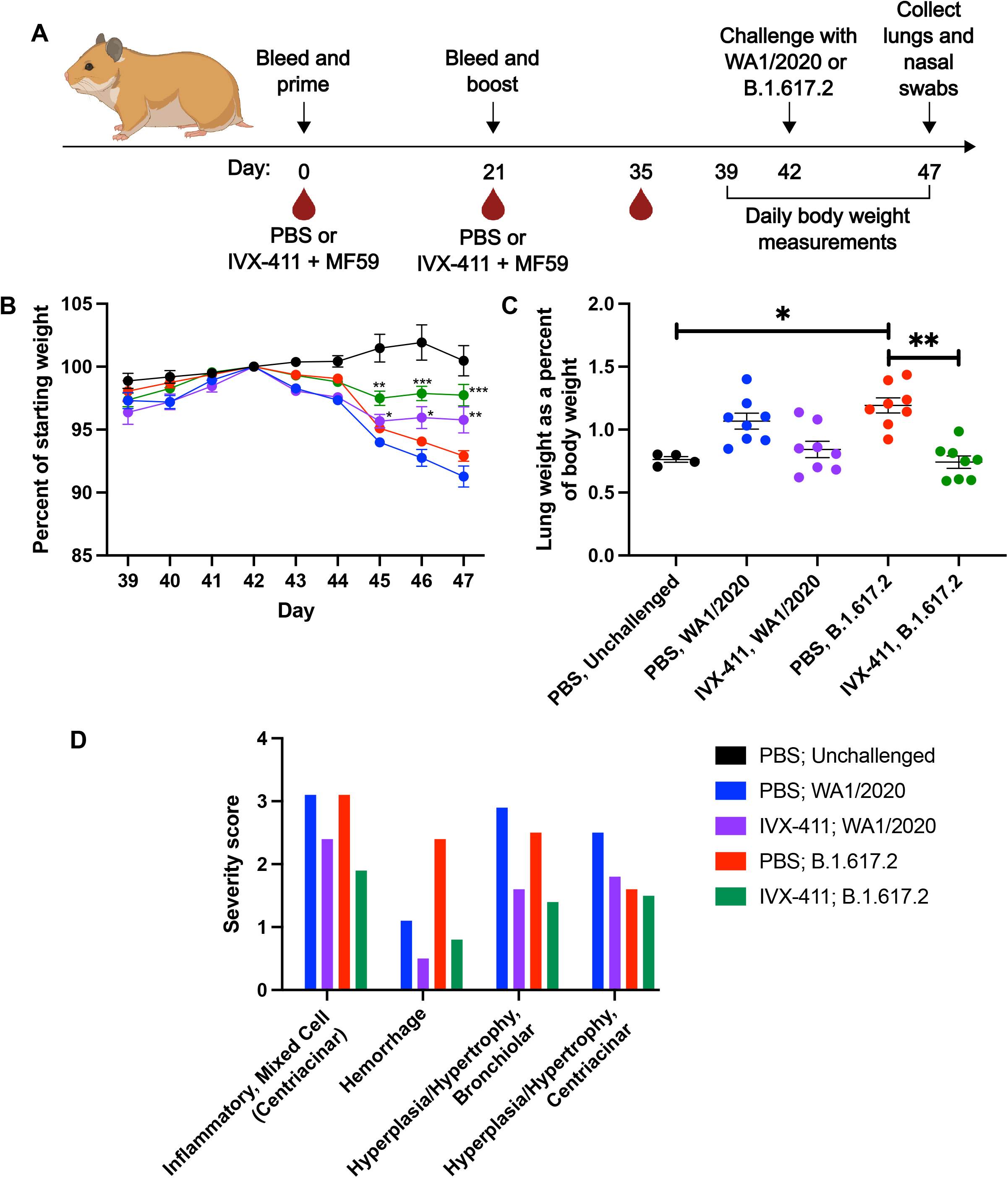
Immunization of Syrian Golden hamsters with IVX-411 reduces severity of SARS-CoV-2 infection. **(A)** Male Syrian Golden hamsters were immunized IM with phosphate buffered saline (PBS) or IVX-411 formulated with MF59 on days 0 and 21. Hamsters in all groups except the PBS, Unchallenged group were inoculated intranasally with one of two SARS-CoV-2 isolates on Day 42. All hamsters were bled on days 0, 21, and 35. Body weights were recorded daily starting on Day 39 until Day 47 when the animals were sacrificed (n = 8). **(B)** Body weights represented as percentage of starting weight prior to challenge. **(C)** Lung weights represented as a percentage of total body weight post challenge. **(D)** Histopathological analysis was performed on lungs collected from each animal. Findings were graded on a scale of 1-5 (1 = Minimal, 2 = Mild, 3 = Moderate, 4 = Marked, 5= Severe). The average severity score for each group is reported. P-values were calculated in GraphPad Prism 9 using a Mann-Whitney test or a Kruskal-Wallis test followed by a Dunn’s multiple comparisons test. *, P< 0.05; **, P<0.01; ***, P<0.001; ****, P<0.0001.

As SARS-CoV-2-induced pneumonia develops, lung weight increases relative to body weight due to cellular infiltrates, edema, and a decrease in overall body weight. Consequently, we investigated the ability of immunization with IVX-411 to prevent an increase in lung weight. As expected, hamsters in the PBS, WA1/2020 and PBS, B.1.617.2 groups had increased lung weight compared to hamsters in the PBS, Unchallenged group **(Figure 4C)**. However, this was only statistically significant in animals challenged with B.1.617.2. Similarly, unimmunized animals challenged with either WA1/2020 or B.1.617.2 had a 1.2 or 1.6-fold increase in lung weight as a percent of body weight respectively compared to their IVX-411 immunized counterparts. Immunization with IVX-411 led to a statistically significant reduction in lung weight in the context of challenge with B.1.617.2 and a trend towards lower lung weights in hamsters challenged with WA1/2020.

In addition to lung and body weight, the impact of IVX-411 immunization on histopathologic changes associated with SARS-CoV-2 infection were also assessed **(Figure 4D)**. The two challenge strains led to slightly different tissue changes in unimmunized animals. B1.617.2 infection led to more hemorrhage and less centriacinar and bronchiolar epithelial hypertrophy/hyperplasia compared to WA1/2020. The impact of IVX-411 immunization on lung pathology was the most notable in animals challenged with B1.617.2. Animals immunized with IVX-411 and then challenged with B1.617.2 had milder inflammation, reduced pulmonary hemorrhage, and decreased severity of bronchiolar epithelial hyperplasia/hypertrophy compared to unimmunized animals. In the case of challenge with WA1/2020, immunization resulted in more subtle decreases in the severity of inflammation, pulmonary hemorrhage, and centriacinar hyperplasia/hypertrophy. The severity of bronchiolar epithelial hyperplasia/hypertrophy was markedly improved when animals challenged with WA1/2020 had been previously immunized with IVX-411.

To further assess the level of protection conferred by immunization with IVX-411, lung tissue and nasal swabs were assessed for genomic RNA (N gene) and subgenomic RNA (E gene) by RT-qPCR. Genomic RNA measures both viable and nonviable viral genomes whereas subgenomic RNA is a measure of replicating virus levels. IVX-411 immunization significantly reduced the number of N gene copies per gram of lung tissue 31 and 66-fold and the number of N gene copies per nasal swab 4.4 and 3-fold in hamsters challenged with WA1/2020 and B.1.617.2, respectively **(Figure 5A)**. Consistent with the N gene results, IVX-411 immunization reduced the number of E gene copies per gram of lung tissue 60 and 132-fold and the number of E gene copies per nasal swab 5 and 3-fold in hamsters challenged with WA1/2020 and B.1.617.2, respectively **(Figure 5B)**. However, the decreases in N and E gene copies in nasal swab samples collected from immunized animals did not reach the level of statistical significance.

**Figure 5:**
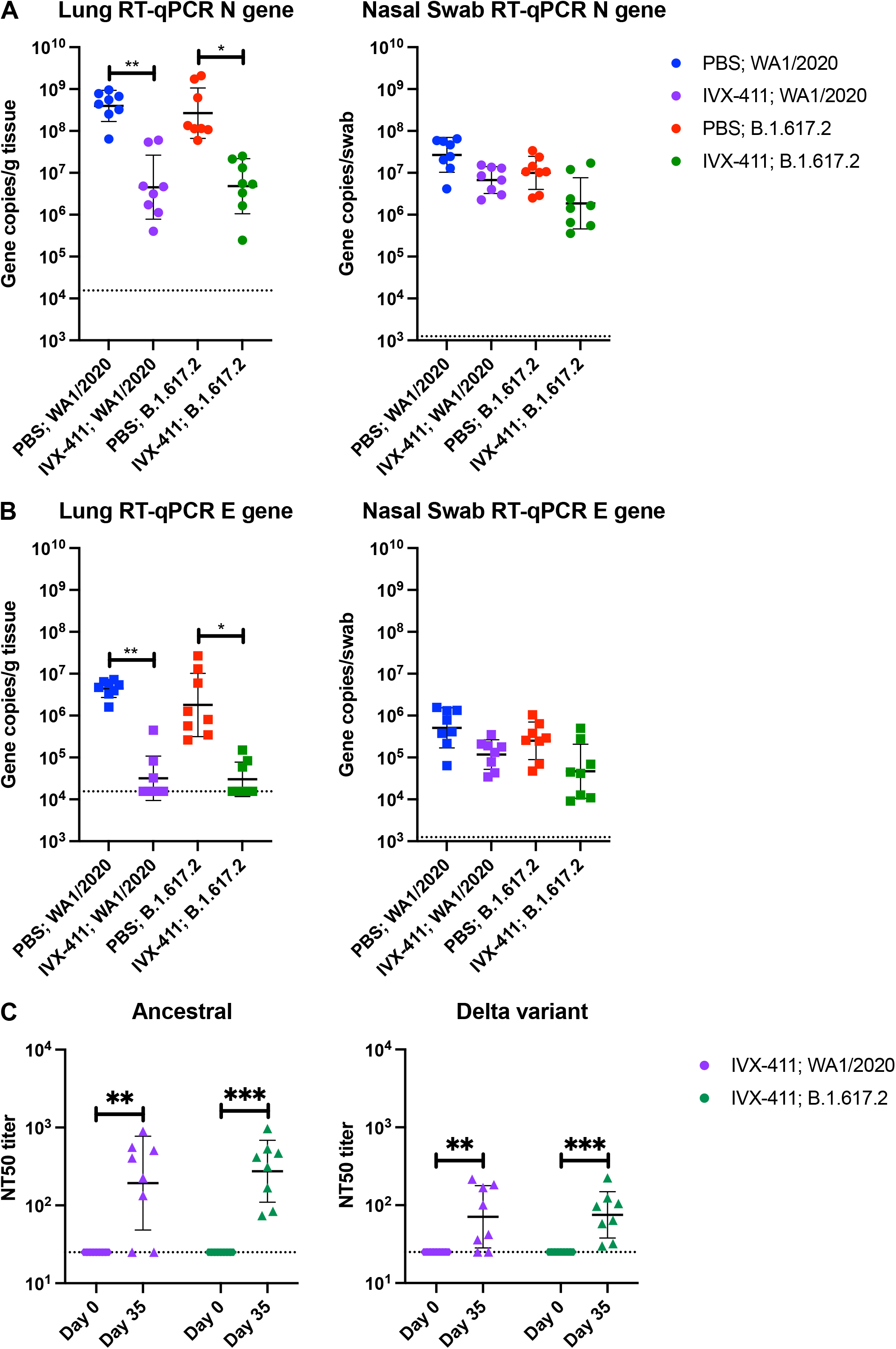
Prophylactic immunization with IVX-411 elicits neutralizing antibodies and results in decreased viral load upon challenge in Syrian Golden hamsters. Number of N **(A)** or E **(B)** gene copies present in lung tissue or nasal swab of Syrian Golden hamsters challenged with WA1/2020 or B.1.617.2 (n = 8). **(C)** Neutralizing antibody titers against the ancestral strain and delta variant on Day 0 and Day 35 in hamsters immunized with IVX-411. Dashed horizontal line represents the lower limit of quantitation. P-values were calculated in GraphPad Prism 9 using a Mann-Whitney test or a Kruskal-Wallis test followed by a Dunn’s multiple comparisons test. *, P< 0.05; **, P<0.01; ***, P<0.001; ****, P<0.0001.

Finally, to determine whether neutralizing antibodies elicited by IVX-411 vaccination were contributing to the observed protection, we evaluated the neutralizing antibody response elicited in the animals immunized with IVX-411 prior to challenge using a cell-based PNA. On Day 35, neutralizing antibody titers against both the ancestral and delta variant were present in six out of eight IVX-411, WA1/2020 hamsters and all eight IVX-411, B.1.617.2 hamsters **(Figure 5C)**. Neutralizing titers against the ancestral strain increased approximately 14.5-fold in IVX-411 immunized animals compared to baseline. A roughly 3.8-fold increase in neutralizing titers against the delta variant was also observed. Overall, these results demonstrate that prophylactic immunization with IVX-411 is protective against severe manifestations of SARS-CoV-2 infection in Syrian Golden hamsters.

## Discussion

As SARS-CoV-2 becomes endemic, there is great interest in identifying vaccines with improved durability, less reactogenicity, and the potential for combination with other vaccines, such as seasonal influenza. VLPs have been shown to be highly immunogenic and possess many of the characteristics that may be of value for a next-generation SARS CoV-2 vaccine. Here, we show that IVX-411, a computationally designed VLP presenting the RBD of the ancestral SARS-CoV-2 strain, is immunogenic in two rodent models. In mice, immunization with IVX-411 elicits neutralizing titers against the ancestral strain as well as three VOCs. These neutralizing antibody titers are durable and higher than those generated in response to immunization with a soluble spike protein. Critically, immunization with IVX-411 is protective in a Syrian Golden hamster challenge model. Together, these results further illustrate the potential of VLPs to be utilized as a modality for future SARS-CoV-2 vaccines.

These data for IVX-411 are consistent with a large number of preclinical studies on I53-50 immunogens displaying RBD antigens from SARS-CoV-2 and/or other sarbecoviruses [24,31,38], and further demonstrate the precision and immunogenic potency of this platform. In naïve animals, higher neutralizing titers against SARS-CoV-2 were observed after immunization with I53-50 VLPs displaying 60 copies of the RBD compared to soluble S protein trimers and trimerized RBD antigens [24,39]. Interestingly, we observed IVX-411 generated higher neutralizing antibody titers in naïve mice than in Syrian Golden hamsters. However, differences in neutralizing antibody titers between different animal models are consistent with what others have reported [32,40]. Further, we observed improved potency in naïve animals by formulating IVX-411 with MF59, an oil-in-water emulsion. Such results have been externally replicated for I53-50-based SARS-CoV-2 vaccines in naïve human patients [38,41–43]. Our results are immunogenically similar to those observed from other RBD-based, single-component, SARS-CoV-2 VLP vaccines, which use nanoparticles such as ferritin [44,45], hepatitis B surface antigen [46], or mi3 (a variant of the I3-01 VLP) [47–50], as well as biochemical conjugation methods such as SpyCatcher and sortase [45,47]. In comparison to these single-component VLP platforms, the two-component nature of I53-50 allows for simplified and modular manufacturing of highly defined immunogen structures through *in vitro* assembly. The precise structure of the final VLP product is enabled both by direct genetic fusion between antigens and CompA, eliminating the need for conjugation, as well as dependable complete and cooperative assembly of the nanoparticle structure due to highly-specific designed interactions between CompA and CompB [35,37]. Finally, the manufacturing and clinical validation of I53-50-based vaccines for SARS-CoV-2 [43] and RSV (Icosavax, Inc. unpublished data) confirms that this platform is scalable, manufacturable, and potently immunogenic while maintaining low reactogenicity, with ample potential to extend to other viral and bacterial indications.

The emergence of the omicron SARS-CoV-2 variant in November, 2021 led to a surge of infections in December, 2021 through March, 2022 [1]. Given the ability of the omicron variant to escape preexisting immunity elicited by both natural infection and immunization, updated booster vaccines targeting the omicron variant as well as the ancestral strain are now recommended for those 12 and older. The I53-50 VLP platform utilized in IVX-411 has the potential to combine multiple SARS-CoV-2 antigens to direct immune responses against several VOCs, including omicron. This platform also has the potency and low reactogenicity needed to generate combination vaccines which could include additional VLP-based vaccines against other viruses. Prior experience with omicron suggests that future updates to SARS-CoV-2 vaccines may be necessary as well as routine booster immunizations. The modular ability for antigen presentation on I53-50 VLPs in response to emerging VOCs, combined with the potential use of high-yield, stabilized RBD designs [39], can enable this platform to meet many needs for future COVID-19 vaccination.

## Declaration of Competing Interest

All authors are employees and stockholders of Icosavax, Inc.

## Acknowledgements

This work was supported in part by the Bill & Melinda Gates Foundation [INV-022092]. Under the grant conditions of the Foundation, a Creative Commons Attribution 4.0 Generic License has already been assigned to the Author Accepted Manuscript version that might arise from this submission. Thanks to Seqirus for providing MF59. Subfigures 2A, 3A, and 4A were created with BioRender.com. Figure 1A was made with UCSF ChimeraX, developed by the Resource for Biocomputing, Visualization, and Informatics at UCSF, with support from NIH R01GM129325 and the Office of Cyber Infrastructure and Computational Biology, NIAID.

## Abbreviations

COVID-19: coronavirus disease 2019
SARS-CoV-2: severe acute respiratory syndrome coronavirus 2
VOCs: variants of concern
S: spike protein
RBD: receptor binding domain
ACE2: angiotensin-converting enzyme 2
VLPs: virus-like particles
LLPCs: long-lived plasma cells
IM: intramuscular
SEC: size exclusion chromatography
DLS: dynamic light scattering
BLI: biolayer interferometry
nsTEM: negative stain transmission electron microscopy
PNA: pseudo-particle neutralization assay

## Figure captions

**Supplemental Figure 1: (A)** UV-Vis spectroscopy and **(B)** SDS-PAGE of purified RBD-CompA trimeric component.

## References

[1] WHO Coronavirus (COVID-19) dashboard n.d. https://covid19.who.int (Accessed July 6, 2022).

[2] Zhou P, Yang X-L, Wang X-G, Hu B, Zhang L, Zhang W, et al. A pneumonia outbreak associated with a new coronavirus of probable bat origin. Nature 2020;579:270–3.

[3] Zhu N, Zhang D, Wang W, Li X, Yang B, Song J, et al. A novel Coronavirus from patients with pneumonia in China, 2019. N Engl J Med 2020;382:727–33.

[4] Mehandru S, Merad M. Pathological sequelae of long-haul COVID. Nat Immunol 2022;23:194–202.

[5] Rogers TF, Zhao F, Huang D, Beutler N, Burns A, He W-T, et al. Isolation of potent SARS-CoV-2 neutralizing antibodies and protection from disease in a small animal model. Science 2020;369:956–63.

[6] Zost SJ, Gilchuk P, Case JB, Binshtein E, Chen RE, Nkolola JP, et al. Potently neutralizing and protective human antibodies against SARS-CoV-2. Nature 2020;584:443–9.

[7] Liu L, Wang P, Nair MS, Yu J, Rapp M, Wang Q, et al. Potent neutralizing antibodies against multiple epitopes on SARS-CoV-2 spike. Nature 2020;584:450–6.

[8] Brouwer PJM, Caniels TG, van der Straten K, Snitselaar JL, Aldon Y, Bangaru S, et al. Potent neutralizing antibodies from COVID-19 patients define multiple targets of vulnerability. Science 2020;369:643–50.

[9] Cao Y, Su B, Guo X, Sun W, Deng Y, Bao L, et al. Potent neutralizing antibodies against SARS-CoV-2 identified by high-throughput single-cell sequencing of convalescent patients’ B cells. Cell 2020;182:73-84.e16.

[10] Ke Z, Oton J, Qu K, Cortese M, Zila V, McKeane L, et al. Structures and distributions of SARS-CoV-2 spike proteins on intact virions. Nature 2020;588:498–502.

[11] Huang Y, Yang C, Xu X-F, Xu W, Liu S-W. Structural and functional properties of SARS-CoV-2 spike protein: potential antivirus drug development for COVID-19. Acta Pharmacol Sin 2020;41:1141–9.

[12] Jia HP, Look DC, Shi L, Hickey M, Pewe L, Netland J, et al. ACE2 receptor expression and severe acute respiratory syndrome coronavirus infection depend on differentiation of human airway epithelia. J Virol 2005;79:14614–21.

[13] Steffen TL, Stone ET, Hassert M, Geerling E, Grimberg BT, Espino AM, et al. The receptor binding domain of SARS-CoV-2 spike is the key target of neutralizing antibody in human polyclonal sera. BioRxiv 2020. https://doi.org/10.1101/2020.08.21.261727.

[14] Piccoli L, Park Y-J, Tortorici MA, Czudnochowski N, Walls AC, Beltramello M, et al. Mapping neutralizing and immunodominant sites on the SARS-CoV-2 spike receptor-binding domain by structure-guided high-resolution serology. Cell 2020;183:1024-1042.e21.

[15] Lai C-C, Ko W-C, Chen C-J, Chen P-Y, Huang Y-C, Lee P-I, et al. COVID-19 vaccines and thrombosis with thrombocytopenia syndrome. Expert Rev Vaccines 2021;20:1027–35.

[16] Ferdinands JM, Rao S, Dixon BE, Mitchell PK, DeSilva MB, Irving SA, et al. Waning 2-dose and 3-dose effectiveness of mRNA vaccines against COVID-19-associated emergency department and urgent care encounters and hospitalizations among adults during periods of Delta and Omicron variant predominance - VISION Network, 10 states, August 2021-January 2022. MMWR Morb Mortal Wkly Rep 2022;71:255–63.

[17] Moreira ED Jr, Kitchin N, Xu X, Dychter SS, Lockhart S, Gurtman A, et al. Safety and efficacy of a third dose of BNT162b2 Covid-19 vaccine. N Engl J Med 2022;386:1910–21.

[18] Menni C, May A, Polidori L, Louca P, Wolf J, Capdevila J, et al. COVID-19 vaccine waning and effectiveness and side-effects of boosters: a prospective community study from the ZOE COVID Study. Lancet Infect Dis 2022;22:1002–10.

[19] Barda N, Dagan N, Cohen C, Hernán MA, Lipsitch M, Kohane IS, et al. Effectiveness of a third dose of the BNT162b2 mRNA COVID-19 vaccine for preventing severe outcomes in Israel: an observational study. Lancet 2021;398:2093–100.

[20] Magen O, Waxman JG, Makov-Assif M, Vered R, Dicker D, Hernán MA, et al. Fourth dose of BNT162b2 mRNA Covid-19 vaccine in a nationwide setting. N Engl J Med 2022;386:1603–14.

[21] Gardner BJ, Kilpatrick AM. Estimates of reduced vaccine effectiveness against hospitalization, infection, transmission and symptomatic disease of a new SARS-CoV-2 variant, Omicron (B.1.1.529), using neutralizing antibody titers. BioRxiv 2021. https://doi.org/10.1101/2021.12.10.21267594.

[22] Lopez Bernal J, Gower C, Andrews N, Public Health England Delta Variant Vaccine Effectiveness Study Group. Effectiveness of covid-19 vaccines against the B.1.617.2 (delta) variant. Reply. N Engl J Med 2021;385:e92.

[23] Higdon MM, Baidya A, Walter KK, Patel MK, Issa H, Espié E, et al. Duration of effectiveness of vaccination against COVID-19 caused by the omicron variant. Lancet Infect Dis 2022. https://doi.org/10.1016/S1473-3099(22)00409-1.

[24] Walls AC, Fiala B, Schäfer A, Wrenn S, Pham MN, Murphy M, et al. Elicitation of potent neutralizing antibody responses by designed protein nanoparticle vaccines for SARS-CoV-2. Cell 2020;183:1367-1382.e17.

[25] Marcandalli J, Fiala B, Ols S, Perotti M, de van der Schueren W, Snijder J, et al. Induction of potent neutralizing antibody responses by a designed protein nanoparticle vaccine for respiratory syncytial virus. Cell 2019;176:1420-1431.e17.

[26] Bachmann MF, Rohrer UH, Kündig TM, Bürki K, Hengartner H, Zinkernagel RM. The influence of antigen organization on B cell responsiveness. Science 1993;262:1448–51.

[27] Wang JW, Roden RBS. Virus-like particles for the prevention of human papillomavirus-associated malignancies. Expert Rev Vaccines 2013;12:129–41.

[28] López-Sagaseta J, Malito E, Rappuoli R, Bottomley MJ. Self-assembling protein nanoparticles in the design of vaccines. Comput Struct Biotechnol J 2016;14:58–68.

[29] Bruce MG, Bruden D, Hurlburt D, Zanis C, Thompson G, Rea L, et al. Antibody levels and protection after hepatitis B vaccine: Results of a 30-year follow-up study and response to a booster dose. J Infect Dis 2016;214:16–22.

[30] Kreimer AR, Herrero R, Sampson JN, Porras C, Lowy DR, Schiller JT, et al. Evidence for single-dose protection by the bivalent HPV vaccine—Review of the Costa Rica HPV vaccine trial and future research studies. Vaccine 2018;36:4774–82.

[31] Walls AC, Miranda MC, Schäfer A, Pham MN, Greaney A, Arunachalam PS, et al. Elicitation of broadly protective sarbecovirus immunity by receptor-binding domain nanoparticle vaccines. Cell 2021;184:5432-5447.e16.

[32] Brouwer PJM, Brinkkemper M, Maisonnasse P, Dereuddre-Bosquet N, Grobben M, Claireaux M, et al. Two-component spike nanoparticle vaccine protects macaques from SARS-CoV-2 infection. Cell 2021;184:1188-1200.e19.

[33] Boyoglu-Barnum S, Ellis D, Gillespie RA, Hutchinson GB, Park Y-J, Moin SM, et al. Quadrivalent influenza nanoparticle vaccines induce broad protection. Nature 2021;592:623–8.

[34] King NP, Bale JB, Sheffler W, McNamara DE, Gonen S, Gonen T, et al. Accurate design of co-assembling multi-component protein nanomaterials. Nature 2014;510:103–8.

[35] Bale JB, Gonen S, Liu Y, Sheffler W, Ellis D, Thomas C, et al. Accurate design of megadalton-scale two-component icosahedral protein complexes. Science 2016;353:389– 94.

[36] Ueda G, Antanasijevic A, Fallas JA, Sheffler W, Copps J, Ellis D, et al. Tailored design of protein nanoparticle scaffolds for multivalent presentation of viral glycoprotein antigens. Elife 2020;9:e57659.

[37] Wargacki AJ, Wörner TP, van de Waterbeemd M, Ellis D, Heck AJR, King NP. Complete and cooperative in vitro assembly of computationally designed self-assembling protein nanomaterials. Nat Commun 2021;12:883.

[38] Arunachalam PS, Walls AC, Golden N, Atyeo C, Fischinger S, Li C, et al. Adjuvanting a subunit COVID-19 vaccine to induce protective immunity. Nature 2021;594:253–8.

[39] Ellis D, Brunette N, Crawford KHD, Walls AC, Pham MN, Chen C, et al. Stabilization of the SARS-CoV-2 Spike receptor-binding domain using deep mutational scanning and structure-based design. Front Immunol 2021;12:710263.

[40] Fluckiger A-C, Ontsouka B, Bozic J, Diress A, Ahmed T, Berthoud T, et al. An enveloped virus-like particle vaccine expressing a stabilized prefusion form of the SARS-CoV-2 spike protein elicits highly potent immunity. Vaccine 2021;39:4988–5001.

[41] Arunachalam PS, Feng Y, Ashraf U, Hu M, Walls AC, Edara VV, et al. Durable protection against the SARS-CoV-2 Omicron variant is induced by an adjuvanted subunit vaccine. Sci Transl Med 2022;14:eabq4130.

[42] Grigoryan L, Lee A, Walls AC, Lai L, Franco B, Arunachalam PS, et al. Adjuvanting a subunit SARS-CoV-2 vaccine with clinically relevant adjuvants induces durable protection in mice. NPJ Vaccines 2022;7:55.

[43] Song JY, Choi WS, Heo JY, Lee JS, Jung DS, Kim S-W, et al. Safety and immunogenicity of a SARS-CoV-2 recombinant protein nanoparticle vaccine (GBP510) adjuvanted with AS03: A randomised, placebo-controlled, observer-blinded phase 1/2 trial. EClinicalMedicine 2022;51:101569.

[44] Joyce MG, Chen W-H, Sankhala RS, Hajduczki A, Thomas PV, Choe M, et al. SARS-CoV-2 ferritin nanoparticle vaccines elicit broad SARS coronavirus immunogenicity. Cell Rep 2021;37:110143.

[45] Saunders KO, Lee E, Parks R, Martinez DR, Li D, Chen H, et al. Neutralizing antibody vaccine for pandemic and pre-emergent coronaviruses. Nature 2021;594:553–9.

[46] Dalvie NC, Tostanoski LH, Rodriguez-Aponte SA, Kaur K, Bajoria S, Kumru OS, et al. SARS-CoV-2 receptor binding domain displayed on HBsAg virus-like particles elicits protective immunity in macaques. Sci Adv 2022;8:eabl6015.

[47] Tan TK, Rijal P, Rahikainen R, Keeble AH, Schimanski L, Hussain S, et al. A COVID-19 vaccine candidate using SpyCatcher multimerization of the SARS-CoV-2 spike protein receptor-binding domain induces potent neutralising antibody responses. Nat Commun 2021;12:542.

[48] Dalvie NC, Rodriguez-Aponte SA, Hartwell BL, Tostanoski LH, Biedermann AM, Crowell LE, et al. Engineered SARS-CoV-2 receptor binding domain improves manufacturability in yeast and immunogenicity in mice. Proc Natl Acad Sci U S A 2021;118:e2106845118.

[49] Cohen AA, Gnanapragasam PNP, Lee YE, Hoffman PR, Ou S, Kakutani LM, et al. Mosaic nanoparticles elicit cross-reactive immune responses to zoonotic coronaviruses in mice. Science 2021;371:735–41.

[50] Hsia Y, Bale JB, Gonen S, Shi D, Sheffler W, Fong KK, et al. Design of a hyperstable 60-subunit protein dodecahedron. [corrected]. Nature 2016;535:136–9.

