## Supplemental Figure 1 for "Virus-like particle displaying SARS-CoV-2 receptor binding domain elicits neutralizing antibodies and is protective in a challenge model"

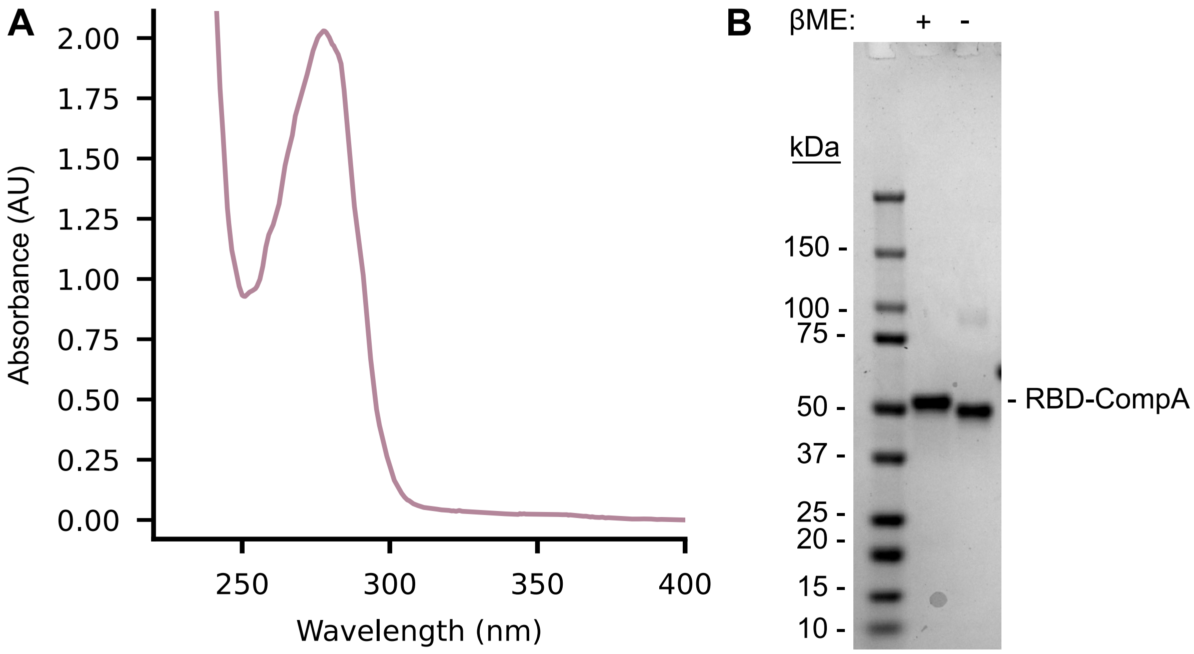


**Supplemental Figure 1: (A)** UV-Vis spectroscopy and **(B)** SDS-PAGE of purified RBD-CompA trimeric component.
